# Unraveling the Dual Immunomodulatory and Immunogenic Roles of the Central Conserved Cysteine-Rich Region in Respiratory Syncytial Virus G Protein

**DOI:** 10.64898/2025.12.06.692521

**Authors:** J Gutman, AL Paletta, F Birnberg-Weiss, CA Prato, A Boudgouste, CJ Goldin, S Sastre, A Byrne, P Pakci, FP Polack, J Dvorkin, A Zeida, MT Caballero, GC Fernández, MV Tribulatti, D Alvarez-Paggi, SA Esperante

## Abstract

Respiratory syncytial virus (RSV) causes severe respiratory disease in infants and high-risk adults, in part by subverting host immunity. The RSV G glycoprotein’s central conserved cysteine-rich domain (CCD) contains a CX3C motif implicated in immune modulation, but the relationship between CCD redox state, structure and function is unresolved. We recombinantly expressed a CCD peptide (Gpep, residues 149–196) and combined structural and biophysical characterizations with cellular immunology and human serology to define how redox-dependent conformations govern immunogenicity and immunomodulation. Reduced Gpep is compact and rapidly folds via a dominant intermediate into an oxidized, extended monomer; at higher concentrations intermolecular disulfide isomerization produces covalent oligomers. Functionally, monomeric Gpep potently suppresses innate and adaptive activation inhibiting LPS- or UV-inactivated RSV-induced maturation of mouse bone marrow-derived dendritic cells, reducing antigen-specific CD4+ T cell proliferation and IFN-γ production, and attenuating multiple human neutrophil responses (chemotaxis, CD11b upregulation, ROS, MPO release and NET formation), without cytotoxicity. Oligomerized Gpep lacks these suppressive activities. To link molecular mechanism and human exposure, analysis of sera from an ambulatory pediatric cohort (0–72 months) showed a progressive transient increase in the anti-F/anti-G IgG ratio with repeated early RSV exposures, maturating into a functional 2 to 3 ratio. This serologic shift is consistent with previously reported enrichment of F-directed neutralizing immunity. We propose a redox-dependent immune-evasion model in which soluble, monomeric G mediates transient immunosuppression that is removed by disulfide-driven oligomerization, which may occur in membrane-bound G. This may impact therapeutic strategies that commonly favor F-focused responses.

**Importance:** Respiratory syncytial virus (RSV) remains a leading cause of severe respiratory disease in young children and high-risk adults. This study identifies a chemical switch in a small conserved region of the RSV attachment protein that changes its shape and immune activity: when the region is present as a soluble, single chain, it transiently suppresses key innate and adaptive immune cells, but this activity is lost upon disulfide-mediated oligomerization. We also show that repetitive RSV exposures in early life bias the antibody response, initially boosting antibody responses toward the Fusion protein, rather than this conserved central region. Together, these results may uncover a mechanism by which RSV shapes the host immune response explaining the features of antibody development in children. Understanding this redox-dependent balance between immune evasion and antigenicity will inform safer vaccine and antibody strategies against RSV.

## Introduction

Respiratory syncytial virus (RSV) is a leading cause of severe lower respiratory tract infections, particularly affecting children under 5 years of age and high-risk populations such as the elderly and immunocompromised individuals^1–4^. Although RSV infection activates the adaptive immune system, the resulting immunity is typically short-lived and non-sterilizing. This leads to recurrent mild RSV infections throughout life and severe disease episodes in vulnerable groups, including immunocompromised individuals and those over 65 years old^5^. Several viral factors alter the host immune microenvironment, promoting disease pathogenesis and recurrent infections^6,7^. Most mechanisms of RSV immune evasion act by suppressing the antiviral interferon response, thereby disrupting both innate and adaptive immunity^8–10^. The RSV genome is a negative-sense, single-stranded RNA encoding 11 proteins from 10 open reading frames (ORFs): NS1, NS2, N, P, M, SH, G, F, M2-1, M2-2, and L11^11^. Among these, the F and G glycoproteins are key surface proteins involved in viral entry, replication, and immune modulation. The F glycoprotein, responsible for fusing the viral and host cell membranes, constitutes the sole antigenic component in recently approved vaccines for older adults and pregnant women^12,13^. Additionally, it is the primary target for prophylactic monoclonal antibodies in preterm and high-risk infants^14^.

The RSV envelope G protein (mG) is a type II transmembrane glycoprotein comprised of a short cytoplasmic N-terminal region (residues 1–37), a transmembrane domain (residues 38–66), and a heavily glycosylated extracellular domain (residues 67–312)^15^. The ectodomain contains two highly variable, intrinsically disordered, and extensively N-and O-glycosylated mucin-like regions, which are implicated in immune evasion^16^. These regions are separated by a central conserved cysteine-rich domain (CCD) and a heparin-binding domain (HBD) responsible for host cell attachment^17^. The CCD is defined by four cysteine residues forming a cysteine noose stabilized by two disulfide bonds (Cys173–Cys186 and Cys176–Cys182 with a 1-4, 2-3 connectivity)^18^. This structure contains the CX3C motif, which interacts with the chemokine receptor CX3CR1^19,20^. An alternative translation initiation at Met48 yields a secreted G ectodomain (sG, residues 67–298) that retains the CCD, HBD, and mucin-like regions. The sG form is proposed to act as an immune decoy by capturing neutralizing antibodies and modulating CX3C chemokine signaling, thereby contributing to pathogenesis^21,22^. While structural differences between sG and mG are not fully elucidated, evidence suggests that mG can form covalently linked dimers through intercatenary disulfide bonds^23^, potentially leading to environment-dependent regulation of CCD structure and function through a redox-based mechanism.

Substantial evidence identifies the CCD as a pivotal virulence factor, facilitating viral entry^19^, modulating host immune responses, and contributing to disease pathogenesis^24^. The CCD, together with HBD-glycosaminoglycan interactions, mediates initial cell surface attachment via the CX3C motif’s interaction with CX3CR1 on human airway epithelial cells^19^. This interaction triggers the production of RANTES, IL-8, and fractalkine, while suppressing IL-15, IL1-RA, and monocyte chemotactic protein-1, thereby impairing the local immune response^20^. The CCD’s CX3C motif reduces antiviral T cell responses, affecting CX3CR1+ T cell trafficking to the lung and diminishing populations of crucial immune cells such as IFN-γ-producing CD8+ T cells and IL-4-producing cells^24^. Furthermore, CCD:CX3CR1 binding on dendritic cells impairs their maturation and lymph node migration, suppressing type I/III interferon and other pro-inflammatory cytokines^8^. The CCD is also a potent inhibitor of the innate immune response during early RSV infection^25^. Disrupting the CX3C:CX3CR1 interaction—either by mutating the CCD or using neutralizing antibodies—restores immune responses^26–29^. In animal models, viruses lacking a functional CX3C motif induce a Th1-biased immune response and higher antibody titers compared to wild-type viruses, with reduced pathology^26,30^.

Paradoxically, the same CCD region is immunogenic and can elicit neutralizing antibody responses. The CX3C motif induces strong antibody production, and anti-G monoclonal antibody therapies have been shown to reduce inflammation and disease severity in RSV-infected models by suppressing Th2-polarized cytokines and chemokines while enhancing interferon responses^31–34^. Thus, the CCD is both immunomodulatory and immunogenic, raising the critical question of whether these seemingly opposing functions overlap at the sequence or structural level. The G protein and particularly its CCD containing the CX3C motif have been extensively investigated as vaccine targets. However, vaccines based on recombinant glycosylated sG have failed to induce robust neutralizing antibody responses and have led to enhanced lung pathology upon RSV challenge, including eosinophilic and neutrophilic infiltration and elevated Th2 cytokines/chemokines^35^. In contrast, unglycosylated sG produced in *E. coli* induced higher titers of neutralizing antibodies and conferred protection without enhanced pathology^36^. Various nanoparticle, microparticle, and vector-based vaccine platforms displaying the CCD have demonstrated protective efficacy in mice^37,38^, yet some CX3C-targeted vaccines have been linked to vaccine-enhanced disease (VED) in animal models. Strategies to mitigate VED include adjuvantation with TLR2 or TLR4 agonists^39,40^ or co-delivery with IL-35^41^. The challenge remains to elicit protective immunity without inducing VED, a complication seen in early vaccine trials with formalin-inactivated RSV^42^. Understanding the molecular determinants of immunomodulation and inflammation is essential for developing safe and effective RSV vaccines^43–45^. Moreover, identifying the factors that steer immune responses toward protective Th1 rather than Th2 phenotypes is critical to overcoming this paradox.

In this study, a peptide (Gpep) comprising the RSV CCD was recombinantly expressed and purified. Its oxidative folding, putative native conformation, and other possible conformations were characterized by uncovering a structural plasticity that may reflect variations between mG and sG. Functional effects were assessed in vitro using mouse bone marrow-derived dendritic cells and splenocyte-derived cells. The results indicated that Gpep suppresses TLR4 agonist-induced dendritic cell activation, inhibits MHC class II-presented OVA peptide-induced lymphocyte activation, and reduces human neutrophil activation under various stimuli. These effects are associated with the redox conformation of the CCD, which may reflect differences in functionality between membrane-bound and soluble G forms. The study aims to clarify the immunomodulatory and immunogenic roles of the RSV G protein CCD, providing information relevant to RSV immunobiology and vaccine development.

## Materials and Methods

### RSV G peptide expression and purification

The central non-glycosylated region (Gpep, amino acids 149–196) of the RSV strain A2 G protein (Uniprot accession No. P03423) was subcloned into a pMalE vector as an Nt-MBP tag fusion protein. A TEV protease recognition site was added between the Nt-MBP and Gpep-Ct sequences. The construct was confirmed by Sanger sequencing and further transformed into *E. coli* C41 (DE3) cells. Transformed bacteria were grown overnight (ON) at 37°C and 220 rpm in LB medium containing 100 µg/mL ampicillin. An ON culture (DO_600nm_ = 3.0) was diluted 1:100 into fresh LB medium and grown for 3 hours until DO_600nm_ reached 0.6. Protein expression was induced by adding 0.25 mM isopropyl β-D-1-thiogalactopyranoside (IPTG), and the culture was grown 16 h at 30°C. Bacterial cells were pelleted at 3500 rpm for 15 minutes at 4°C. The resulting pellet was resuspended in lysis buffer (25 mM Tris.HCl, pH 7.6, 200 mM NaCl). Resuspended cells were lysed by sonication and centrifuged at 20,000 rpm for 20 minutes at 4°C. The soluble protein was precipitated by adding ammonium sulfate to a final concentration of 80% with continuous stirring at 4°C overnight. The precipitate was collected by centrifugation at 12,000 rpm for 20 minutes at 4°C and resuspended in 50 mL/Litre of culture in amylose buffer A: 20 mM Tris-HCl, pH 7.6, 200 mM NaCl. The resuspended sample was loaded onto an amylose affinity purification column (Cytiva) equilibrated in buffer A. The column was washed with 10 CV of buffer A, and protein elution was performed by adding 20 mM maltose in buffer A. The purified fusion protein was incubated with TEV protease at a 1:100 mass ratio for 16 h at 20°C to cleave Gpep from MBP. Considering the distinct isoelectric points of the two proteins (9.5 for Gpep and 5.1 for MBP), the cleaved Gpep was purified using cation exchange chromatography on a Capto S column (Cytiva). The column was equilibrated with buffer A (50 mM sodium phosphate, pH 7.6). The protein sample was previously dialyzed in buffer A and applied to the column. After washing with 5 CV of buffer A, elution was performed by a 10 CV gradient from 0 to 500 mM NaCl. The Gpep was further purified by size exclusion chromatography in a Superdex 75 column (Cytiva), previously equilibrated in phosphate-buffered saline (PBS, Gibco). Fractions containing the protein of interest were collected and purity was analyzed by Tris-Tricine Gel electrophoresis stained by coomassie-blue and RP-HPLC in a C4 column (see below). Protein concentration was determined spectrophotometrically by using an extinction coefficient of 5500 M^-1^cm^-1^.

### RP-HPLC

The protein sample was analyzed by reverse-phase high-performance liquid chromatography (RP-HPLC) using a C4 column. Separation was achieved with a linear gradient of acetonitrile from 20% to 40% in the presence of 0.1% trifluoroacetic acid (TFA) over 50 minutes at a flow rate of 1 mL/min. Elution was monitored by UV absorbance at 280 nm.

### Thiol quantification

Free thiols in the native or reduced protein sample were quantified using 5,5′-dithiobis(2-nitrobenzoate) (DTNB^46^), which was purchased from Sigma Aldrich and used following the manufacturer’s protocol. Sample was reduced via incubation with 1 mM DTT for one hour, and then desalted with a PD10 column. Calibration was performed with reduced glutathione (GSH, one thiol/molecule)

### Oxidative Folding

The peptide at 1.6 mg/mL was reduced and denatured by incubating for 16 h at 20°C in 50 mM TrisHCl pH 8.0 containing 100 mM DTT and 6.0 M guanidinium chloride. The sample was buffer exchanged in a PD-10 column in 50 mM TrisHCl pH 8.0. The folding reaction was allowed to proceed under four different conditions: A (Control-): 50 mM TrisHCl pH 8.0; B (Control +) 50 mM TrisHCl pH 8.0 containing 0.25 mM β-mercaptoethanol as a thiol initiator, C (GSSG): 50 mM TrisHCl pH 8.0 containing 1.0 mM GSSG or D: redox buffer (GSSG:GSH mixture at a 1:0.5 mM ratio). The folding reaction was stopped by acidification with 1% TFA at different time points, and folding intermediates were analyzed by RP-HPLC.

### Fluorescence spectroscopy

Tryptophan emission fluorescence of Gpep was measured on a fluorescence spectrophotometer (Jasco FP-8500) at 20°C. The spectra were taken in the 310 to 450 nm interval using a 280 nm excitation wavelength (5 nm excitation and emission slits, 0.1 s averaging time). To evaluate disulfide reduction and peptide denaturation probed by its single Trp residue, samples of Gpep at 20 uM were incubated for 16 h at 20°C in 20 mM Tris.HCl pH 8.0 containing the following additives: a) no additive; b) 10 mM DTT; c) 6.0 M guanidinium chloride or d) 10 mM DTT and 6.0 M guanidinium chloride. The experiments were performed in duplicates twice.

### Western Blot

Protein samples were separated by SDS–PAGE under reducing conditions and transferred onto PVDF membranes (Millipore). Membranes were blocked for 1 h at room temperature (RT) with 1% BSA in TBS-0.1% Tween 20 (TBS-T). After washing, membranes were incubated overnight at 4 °C with a rabbit polyclonal anti-Gpep antibody (Sino Biological) diluted in blocking buffer. Following three washes with TBS-T, membranes were incubated for 1 h at RT with IRDye® 680RD goat anti-rabbit IgG (LI-COR) diluted in blocking buffer. Fluorescent signals were detected using the Odyssey Imaging System (LI-COR) according to the manufacturer’s instructions.

### Molecular Dynamics Protocol

Both oxidized and reduced initial atomic coordinates of Gpep were generated from the PDBid 6BLI^47^. After removing crystallographic waters and antibody fragments, hydrogen atoms were added by using the *tleap* module of the AMBER18 suite^48^, assigning protonation states consistent with physiological pH. The initial coordinates were solvated in a truncated-octahedral TIP3P water box extending 12 Å beyond the protein. Counterions were added to neutralize the total charge. Energy minimization and molecular dynamics were performed using the ff14SB force field^49^. Briefly, a standard MD protocol was employed: a two-step minimization, followed by a 0.5 ns NVT equilibration in which the system was heated to 300 K using the Berendsen thermostat, and subsequently switched to NPT conditions to equilibrate the density to ∼1 g·cm^-3^ using the Monte Carlo barostat. A 10 Å cutoff was used for non-bonded interactions, and long-range electrostatics were treated under periodic boundary conditions (PBC) with the particle-mesh Ewald (PME) procedure using a grid spacing of 1 Å^50^. The SHAKE algorithm was applied to constrain bonds to hydrogen atoms, and a 2 fs integration time step was used^51^. After 100 ns of conventional MD, 5 replicas of 200 ns long accelerated molecular dynamics (aMD) simulations were performed to enhance sampling, following the dual-boost scheme^52^. In all cases, the potential-energy boost did not exceed 5% of the total potential-energy or dihedral contributions. The measured properties of interest were reweighted by approximating the total ΔV using a 10th-order Maclaurin series^53^. All simulations were carried out with the AMBER18 package^48^, and visualization and molecular graphics were generated using VMD 1.9.1^54^.

### Detoxification

The purified protein was treated with Pierce High Capacity Endotoxin Removal Resin (Thermo Scientific) according to the manufacturer’s protocol.

### Mice

BALB/cJ, C57BL/6J, and OT-II [B6.Cg-Tg(TcraTcrb)425Cbn/J] breeding pairs were obtained from The Jackson Laboratory (Bar Harbor, ME) and bred in our facilities. The Ethics Committee Board of the Universidad Nacional de San Martín approved all procedures involving animals.

### Immunizations

6–8-week-old female BALB/cJ mice (n=5 per group) were subcutaneously vaccinated with 2 µg of Gpep or control antigen (flagellin) in PBS, in the presence or absence of Alum (50% V/V). Immunizations were administered on day 0, followed by blood collection on day 14. A booster dose was given on day 21, with another blood collection on day 35. The third immunization was performed on day 42, and the final blood sample was taken on day 52.

### Bone Marrow Derived Dendritic Cells (BMDCs) activation assay

Bone marrow was obtained from femurs and tibias of 6- to 10-week-old C57BL/6J male mice by flushing RPMI 1640 medium through the bone interior. The cell suspension was washed, and red blood cells were lysed with red blood cell lysis buffer (Sigma-Aldrich). Cells were cultured at 37°C and 5% CO2 in six-well plates (1.5 × 10^6 cells per well) in 2 mL RPMI 1640 containing 10% FBS, 50 µg/mL gentamicin (Sigma-Aldrich), and 20 ng/mL of recombinant GM-CSF (GenScript). Media was renewed on days 3 and 5, then cells were harvested on day 8, and the expression of DC (CD11c, MHC-II) and activation markers were analyzed by flow cytometry. BMDCs were treated for 18 hours with 500 ng/mL LPS and/or 250 nM, 500 nM, or 1000 nM Gpep. The expression of costimulatory molecules CD80, CD86 and MHCII was analyzed by flow cytometry.

### Costimulation assays

For costimulatory assays, splenocytes (3×10^5^ cells) from OTII mice were cultured for 48 h at 37°C in 5% CO_2_ in flat -bottom, 96 -well plates in 0.2 ml RPMI 1640 medium (Thermo Fisher Scientific, Waltham, MA, USA), in the presence of 10% FBS ( Gibco, Thermo Fisher Scientific, Massachusetts, USA), 2 mM glutamine, and 5 mg/ml Gentamicin (complete medium), in the presence of the cognate OVA_323–339_ peptide (Sigma-Genosys, The Woodlands, TX, USA) and LPS (Sigma) at the indicated concentrations. To assess proliferation, 1 μCi [3H] methylthymidine (New England Nuclear, Newton, MA, USA) was added to each well 18 h before harvesting. For IFN-γ quantification, supernatants were collected 24 hours after stimulation, and IFN-γ was measured using Mouse IFN-γ ELISA Kit, (Biolegend). All assays were performed in quadruplicate.

### Flow Cytometry

Flow Max Cytometer PASIII (Partec, Münster, Germany) and FlowJo software (FlowJo, Ashland, OR, USA) were used.

### Adult blood samples and ethical statement

Blood samples were obtained from healthy volunteer donors who had not taken any medication for at least 10 days before the day of sampling. Blood was obtained by venipuncture of the forearm vein and was drawn directly into plastic tubes containing 3.8 % sodium citrate (Merck). This study was performed according to institutional guidelines (National Academy of Medicine, Buenos Aires, Argentina) and received the approval of the institutional ethics committee (N◦ 10/24/CEIANM), and written informed consent was provided by all the subjects.

### Polymorphonuclear neutrophil isolation

Polymorphonuclear neutrophils (PMN) were isolated by Ficoll-Hypaque gradient centrifugation (Ficoll Pharmacia, Uppsala; Hypaque, Wintthrop Products, Buenos Aires, Argentina) and dextran sedimentation, as previously described^55^. Contaminating erythrocytes were removed by hypotonic lysis. After washing, the cells (96% neutrophils on May Grünwald/Giemsa-stained Cyto-preps) were suspended in RPMI 1640 supplemented with 2 % heat-inactivated fetal calf serum (FCS) and used immediately after.

### Chemotaxis assay

Chemotaxis was quantified using a modification of the Boyden chamber technique^56^. A cell suspension (50 μl) containing 2×10^6^ cells/ml in RPMI with 2 % FCS, was placed in the top wells of a 48-well micro-chemotaxis chamber. A PVP-free polycarbonate membrane (3 μm pore size; Neuro Probe Inc. Gaithersburg MD, USA) separated the cells from lower wells containing either RPMI or the chemotactic stimulus (Gpep 1 µM or fMLP 10^-7^ M). The chamber was incubated for 30 min at 37 °C in a 5% CO_2_ humidified atmosphere. After incubation, the filter was stained with TINCION-15 (Biopur SRL, Rosario, Argentina), and the number of neutrophils on the undersurface of the filter was counted in five random high-power fields (HPF) ×400 for each of triplicate.

### CD11b expression

Neutrophils (5×10^5^) were incubated in the presence of Gpep 1 µM, LPS (0.5 ng/ml) or both for 30 min at 37 °C in 5 % CO_2_. Immediately after, cells were incubated with a specific mouse anti-human CD11b antibody conjugated with phycoerythin (PE) (Dako, Santa Clara, CA, USA). Debris was excluded by FSC-SSC and CD11b expression was analyzed within the gated-viable neutrophils. Mean Fluorescence Intensity (MFI) of the CD11b was determined on 50.000 events.

### ROS production

ROS production was measured by flow cytometry using dihydrorhodamine-123 (DHR, Sigma-Aldrich). Briefly, isolated neutrophils (5×10^5^) were incubated 15 min at 37 °C with 1 µM DHR. Subsequently, cells were incubated with Gpep 1 µM, fMLP 10^-7^ M or both for 30 minutes at 37 °C in 5 % CO_2_. Immediately after, the green fluorescence was determined by flow cytometry.

### Degranulation assay

To evaluate primary o azurophilic granule mobilization, neutrophils (1×10^6^) were incubated in a 24-well plate with Gpep 1 µM, PMA 20 nM (Merck) or both for 2 h at 37 °C in 5 % CO_2_. The MPO release was evaluated in cell-free supernatants by measuring its activity. MPO activity was determined using the specific substrate 3, 3′, 5, 5′-tetramethyl-benzidine (TMB) (Invitrogen, Thermo Fisher) and measuring the resultant absorbance at 450 nm, subtracting the absorbance at 570 nm (450–570 nm).

### NETs formation

Neutrophils (1×10^6^) were incubated in a 24-well plate in the presence of Gpep 1 µM, PMA 20 nM or both for 3 h at 37 °C in 5 % CO_2_. After the incubation period, micrococcal Nuclease S7 (4 U, Roche Diagnostics) was added for 15 min at 37 °C in order to cut and release the DNA associated with NETs from the cell body of neutrophils. Then, inactivation of the enzyme was performed by the addition of 5 mM of EDTA. Supernatants were collected and centrifuged twice, first at 900 xg and then at 9600 xg. To determine the presence of double stranded (ds) DNA, a commercial kit was used (Invitrogen, Thermo Fisher), containing picogreen as the DNA intercalator, which was read in a fluorimeter (DeNovix DS-11 spectrophotometer).

### Cell viability

Neutrophils (5×10^5^) were incubated with Gpep or a positive control of death (PMA, 100 nM) for 1 or 4 h at 37 °C in 5 % CO_2_. After the incubation period, cells were washed and incubated with 7-Amino-Actimycin D (7-AAD; BD Biosciences, 10 µg/ml) for 10 min on ice in the darkness. Immediately after, cell viability was determined by flow cytometry, since 7-AAD is excluded by viable cells but can penetrate cell membranes of dying or dead cells.

### Pediatric blood samples

Peripheral blood samples were obtained from pediatric outpatients attending Hospital de Niños “Sor María Ludovica” (La Plata, Buenos Aires, Argentina). Participants were clinically healthy children between 0 and 72 months of age, with no evidence of acute infection at the time of sampling or during the past month. Blood was collected by venipuncture during routine clinical procedures, and serum was separated and stored at –80 °C until use. The institutional review board of the hospital approved the study. Informed consent for serum sample collection was obtained from all participating parents or legal guardians.

### Determination of IgG Antibodies Against RSV G or F Proteins by ELISA

Indirect ELISAs were performed using flat-bottom high-binding ELISA plates (Costar, Corning, NY, USA) coated overnight at 4 °C with 50 ng per well of recombinant RSV G or RSV F proteins (11070-V08H and 11049-V08B, Sinobiological). Plates were washed three times with washing buffer (TBS + 0.1% Tween-20) and blocked for 1 hour at room temperature (RT) with 1% BSA in TBS (Gibco). After discarding the blocking solution, serial dilutions of sera (human or mouse) were added and incubated for 2 hours at RT. Plates were then washed three times and incubated with HRP-conjugated secondary antibody appropriate for the species tested: mouse anti-human IgG for human sera, goat anti-mouse IgG for mouse sera (ThermoFisher, Waltham, MA, USA) for 1 hour at RT. Following three additional washes, plates were developed with 1-Step Ultra TMB Substrate (ThermoFisher) and the reaction was stopped with 1 N sulfuric acid. Absorbance was measured at 450 nm using a microplate reader (BioTek, Winooski, VT, USA). Antibody titers were calculated by fitting the absorbance values to a four-parameter logistic (4PL) sigmoidal curve using nonlinear regression. The titer was defined as the reciprocal of the serum dilution corresponding to a cutoff value equal to 1.5 times the absorbance of the blank for each plate.

## Results

### 1. The RSV G Central Cystein-Rich Peptide Exhibits Redox-Dependent Conformational Plasticity and Oligomerization

All the accumulated evidence in RSV G points towards the CCD as the critical sequence concentrating immunomodulatory properties, redox-sensitive structural determinants and neutralizing epitopes. Thus, the central cysteine-rich region of the RSV G protein (Gpep, residues 149 to 196) containing the CX3C chemokine motif was recombinantly expressed and purified to high homogeneity (**Fig. Sup 1**). While structural evidence shows that this region adopts a cysteine noose stabilized by two intrachain disulfide bonds (Cys1-Cys4, Cys2-Cys3). (**Fig. Sup 1**), recent experimental evidence in the literature demonstrates that multimeric forms of membrane-associated G (mG) can arise, possibly through intermolecular disulfide bonds, indicating additional redox states beyond the classic cysteine noose.^23^

To systematically probe the redox landscape of Gpep, we characterized its native, fully reduced and fully oxidized states. RP-HPLC (**Fig. 2a)** and thiol quantification using DTNB (Ellman’s reagent for free thiol quantifying^46^, data not shown) confirmed that the recombinant peptide predominantly adopts an oxidized conformation with minimal free thiols, consistent with intrachain formation. Refolding assays of the reduced Gpep under various redox conditions (**Fig. 3**) revealed that the peptide efficiently forms the oxidized state, with evidence of a kinetically trapped intermediate prone to structural rearrangement through thiol-disulfide exchange. Folding rates and intermediate accumulation varied depending on the redox environment, but all conditions ultimately yielded a product matching the recombinant oxidized form.

**Figure 1:**
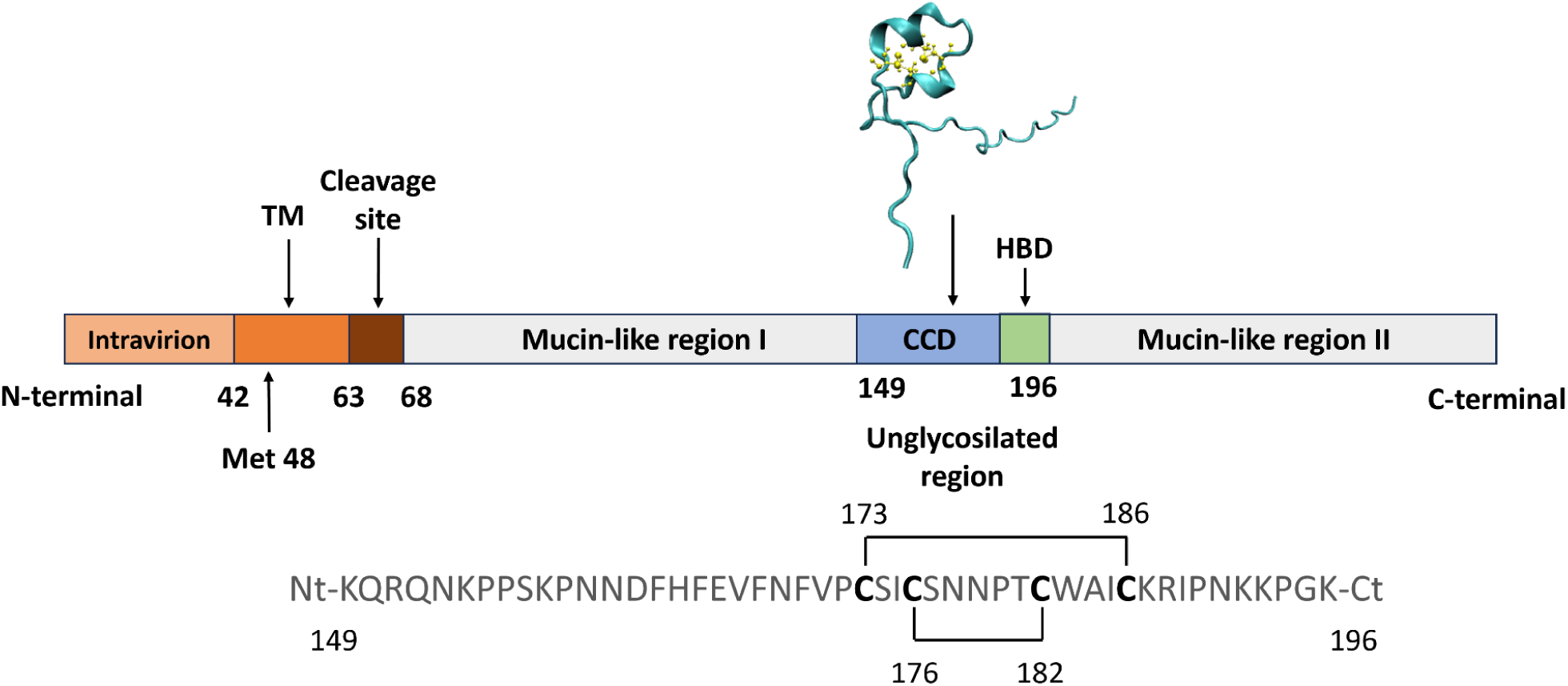
Schematic Representation of the RSV G Protein Structure. The diagram illustrates the regions and functional domains of the RSV G protein, including its cytoplasmic tail, transmembrane domain, cleavage site and the two mucin-like regions (I and II). The central conserved domain (CCD) and the heparin-binding domain (HBD) together constitute the unglycosylated region corresponding to the recombinant peptide Gpep analyzed in this work. The Gpep sequence with its disulfide bond connectivity is shown below.

**Figure 2.**
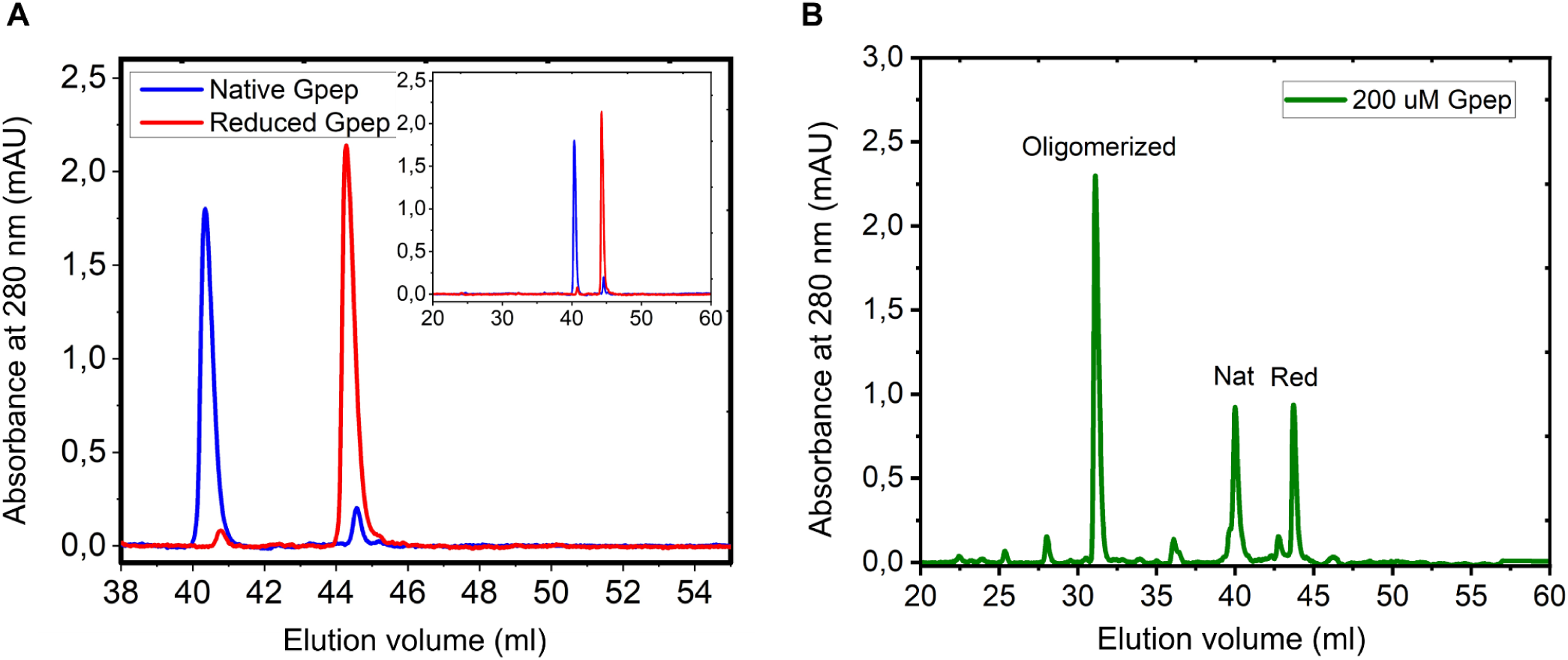
RP-HPLC characterization of reduced, native and oligomeric Gpep. (A) RP-HPLC profile of the native peptide (60 μM, blue trace: elution volume at 40 mL) and its reduced form (60 uM, red trace: elution volume at 44 mL). (B) RP-HPLC profile of Gpep at 200 μM. The major species eluded at 31 mL, corresponding to the oligomerized form.

**Figure 3.**
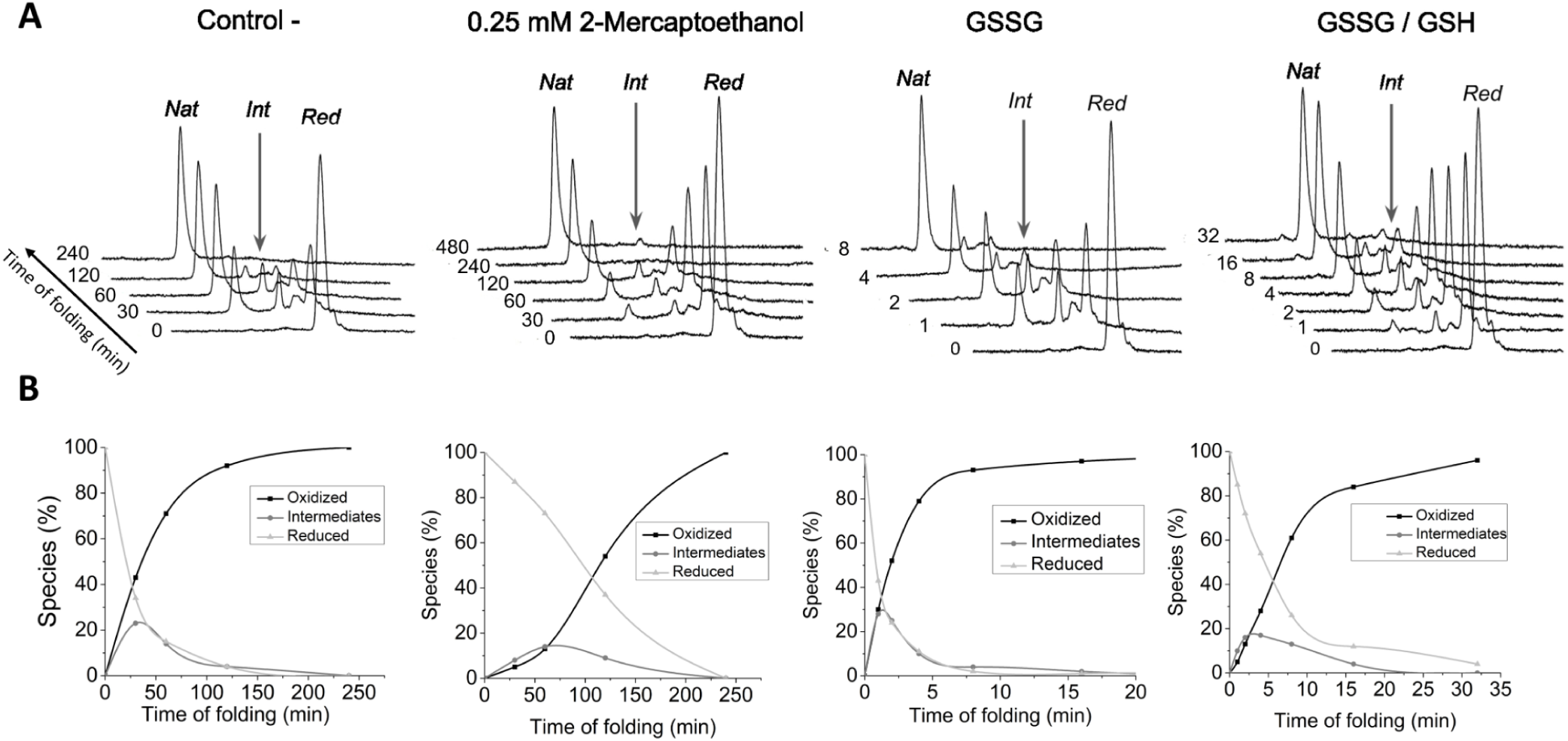
Analysis of the Gpep oxidative folding pathway by RP-HPLC under different redox conditions. RP-HPLC chromatographic profiles of oxidative folding species of Gpep under different redox conditions. Folding was carried out in Tris-HCl buffer (pH 8.0) either without additives (Control −) or in the presence of 0.25 mM 2-mercaptoethanol, 0.5 mM oxidized glutathione (GSSG), or a 0.5 mM GSSG + 1 mM GSH mixture. (A) Chromatographic profiles at different time points. Native (Nat), reduced (Red) and major intermediate (Int) species of Gpep are indicated. (B) Relative abundance of reduced, intermediate, and oxidized species quantified by peak area of RP-HPLC chromatograms shown in A.

Hydrodynamic (**Fig. 4a)** and structural analyses (SEC, SDS-PAGE, and fluorescence spectroscopy) (**Fig. 4b**) revealed that the oxidized Gpep behaves as an elongated monomer in solution, but reduction leads to a partially compact, flexible conformation. Molecular dynamics simulations (**Fig. 4c-e)** further supported the more compact behavior of the reduced Gpep, that could not only be explained by the end-to-end distances observed. This suggests the involvement of other stabilizing elements in the reduced form.

**Figure 4.**
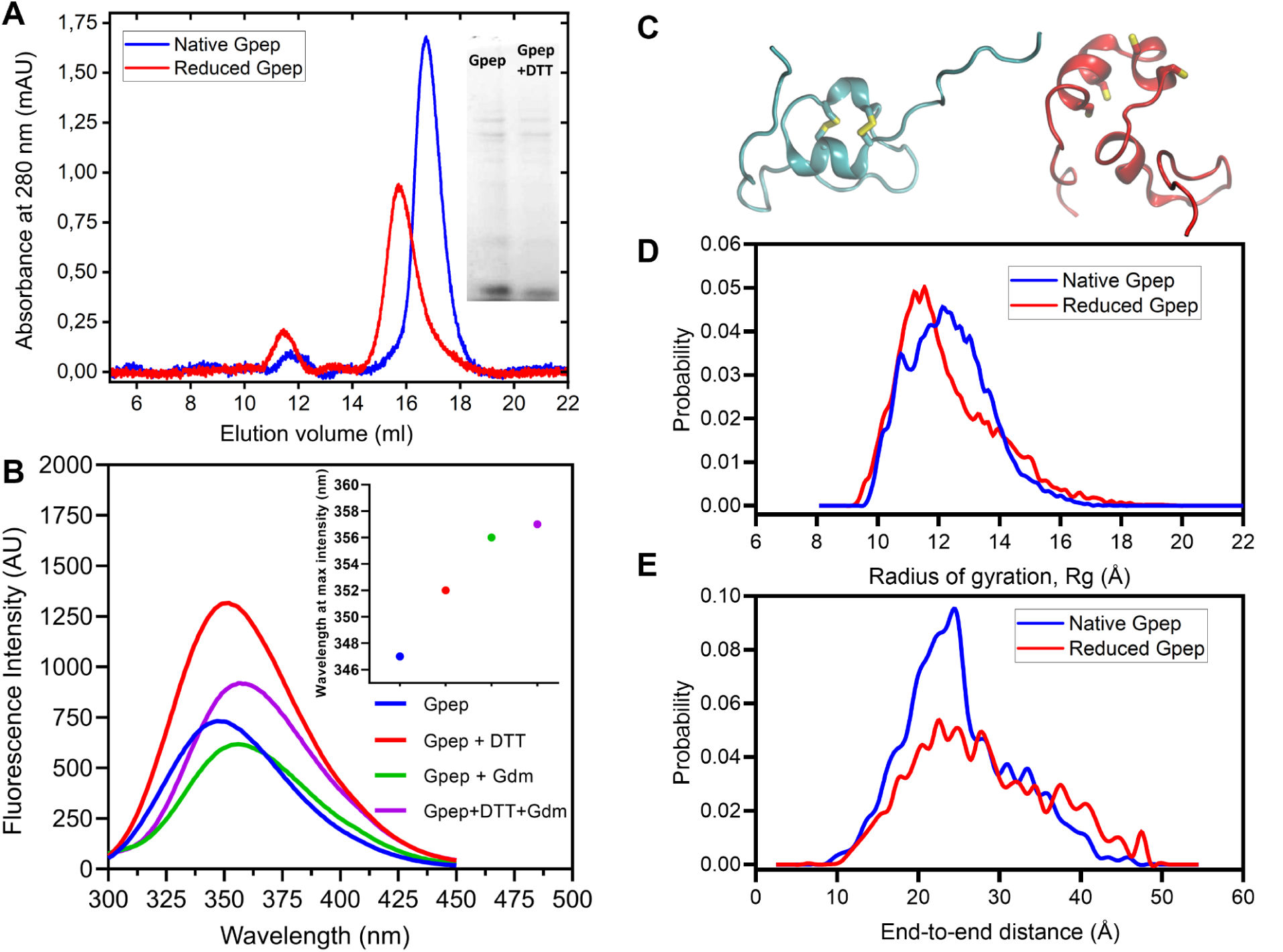
Native Gpep adopts an extended monomeric conformation in solution. (A) Size-exclusion chromatography of native and reduced Gpep on a S75 10/300 column equilibrated in PBS at 20 °C. According to the column calibration, the apparent molecular mass of native Gpep is ∼13.6 kDa and ∼16.2 kDa for reduced Gpep. The globular proteins used for column calibration are: BSA (66.5 kDa), MBP (42 kDa), mCherry (29 kDa) and lysozyme (14.6 kDa). Their elution volumes are indicated by an arrow. (B) Tryptophan fluorescence emission spectra of 20 µM Gpep treated with: 10 mM DTT (red line), 6.0M Gdm·Cl (green line), both reagents (violet line) or no additives (blue lines). Inset: Wavelength of maximum emission of each spectrum. (C) Representative snapshots of both oxidized (blue) and reduced (red) Gpep models obtained by aMD simulations. The four Cys residues are highlighted. (D) Probability distributions of Gpep radius of gyration (*R_g_*, Å) and (E) end-to-end distances (Å).

Importantly, when Gpep was concentrated, we observed a shift in the RP-HPLC profiles coinciding with the appearance of higher molecular weight, disulfide-linked oligomers (**Fig. 2b)**. These oligomeric species were sensitive to reducing agents, indicating that intermolecular disulfide bonds drive their formation. This concentration-dependent oligomerization suggests a structural adaptability of the G region, possibly mirroring the differences observed between the soluble (sG) and membrane-bound (mG) forms of the RSV G protein. It must be stressed that this transition is observed only by changing the peptide concentration without any additional redox stimulus. Such plasticity and redox-dependent behavior may have functional implications for RSV biology.

### 2. The recombinant RSV G-peptide exhibits both immunogenic and immunosuppressive properties

To evaluate the immunogenic potential of the recombinant G central cysteine-rich domain (Gpep), we assessed its ability to present native-like epitopes and elicit functional antibodies. Enzyme-linked immunosorbent assays (ELISA) using sera from healthy adults with prior RSV exposure demonstrated robust recognition of recombinant Gpep, with antibody titers comparable to those against full-length, glycosylated G protein expressed in eukaryotic cells (**Fig- Sup 2a**) This finding substantiates that Gpep preserves native conformational epitopes reflective of the viral protein. Furthermore, sera from BALB/Cj mice immunized with adjuvanted Gpep generated antibodies that specifically recognized the full-length G protein in ELISA (**Fig. Sup 2c**), unequivocally validating that the recombinant peptide presents authentic antigenic determinants. Recognition of Gpep by a commercial anti-G antibody in a Western blot confirms the preservation of its native epitopes (**Fig. Sup 2b**).

Beyond its immunogenicity, the CCD exhibits significant immunomodulatory activity. In mouse bone marrow-derived dendritic cells (BMDCs), recombinant Gpep consistently suppressed activation in response to both LPS and UV-inactivated RSV, as measured by downregulation of surface activation markers CD80, CD86, and MHC-II (**Fig. 5a, 5b**). This indicates that Gpep can modulate innate immune activation under both artificial and physiologically relevant conditions. In ex-vivo assays with splenocytes from OT-II transgenic mice, Gpep inhibited CD4+ T cell proliferation and IFN-gamma production in a concentration dependent manner, as determined by [3H]-thymidine incorporation and cytokine quantification (**Fig. 6a, 6b**). These results demonstrate that Gpep attenuates antigenic specific T cell responses, likely via its effects on dendritic cell function.

**Fig 5:**
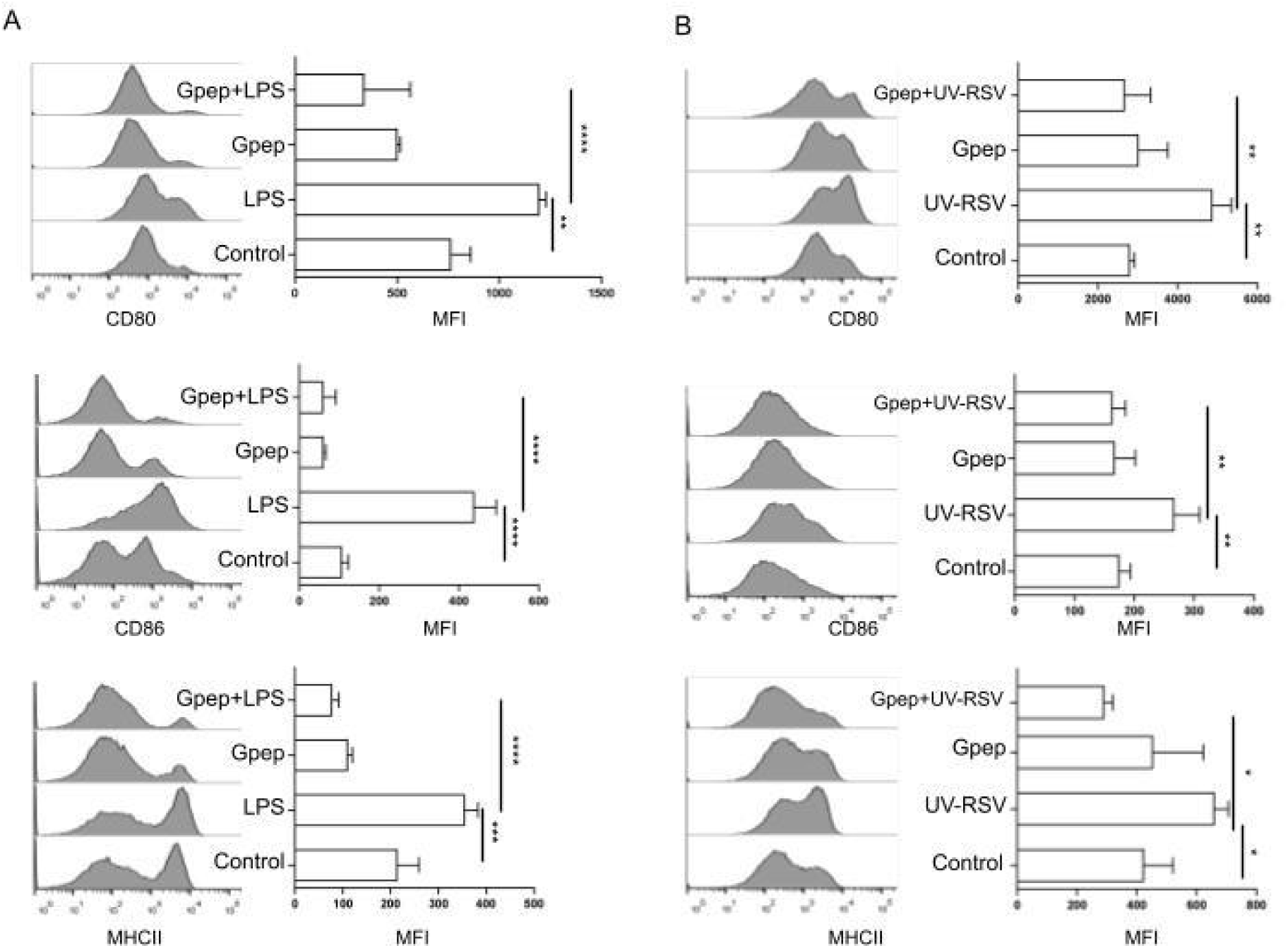
Gpep modulates BMDCs Activation Induced by LPS and RSV. (A). BMDCs were incubated for 18 hours at 37°C in the presence of 500 ng/ml LPS, 1 µM Gpep, or 500 ng/ml LPS plus 1 µM Gpep. Controls are cells without any stimulus. Then, cells were labeled with monoclonal antibodies (mAbs) specific for CD11c, CD80, CD86 and MHCII, and activation was analyzed by flow cytometry in CD11c+ cells. (B). BMDCs were incubated for 18 hours at 37°C in the presence of UV-inactivated RSV (UV-RSV) at a MOI of 500, 1 µM Gpep, or UV-RSV at MOI of 500 plus 1 µM Gpep. Control cells were incubated with RSV inactivation media. Then, cells were labeled with monoclonal antibodies (mAbs) specific for CD11c, CD80, CD86 and MHCII, and activation was analyzed by flow cytometry. Representative histograms depict fluorescence intensity corresponding to the expression of CD80, CD86, and MHCII molecules on the CD11c^+^ gated population. Bars represent the geometric mean fluorescence intensity (MFI) ± SD. *p<0.05, **p<0.01, ***p<0.001, ****p<0.0001 (ANOVA). Depicted assays are representative of at least two independent experiments.

**Fig. 6:**
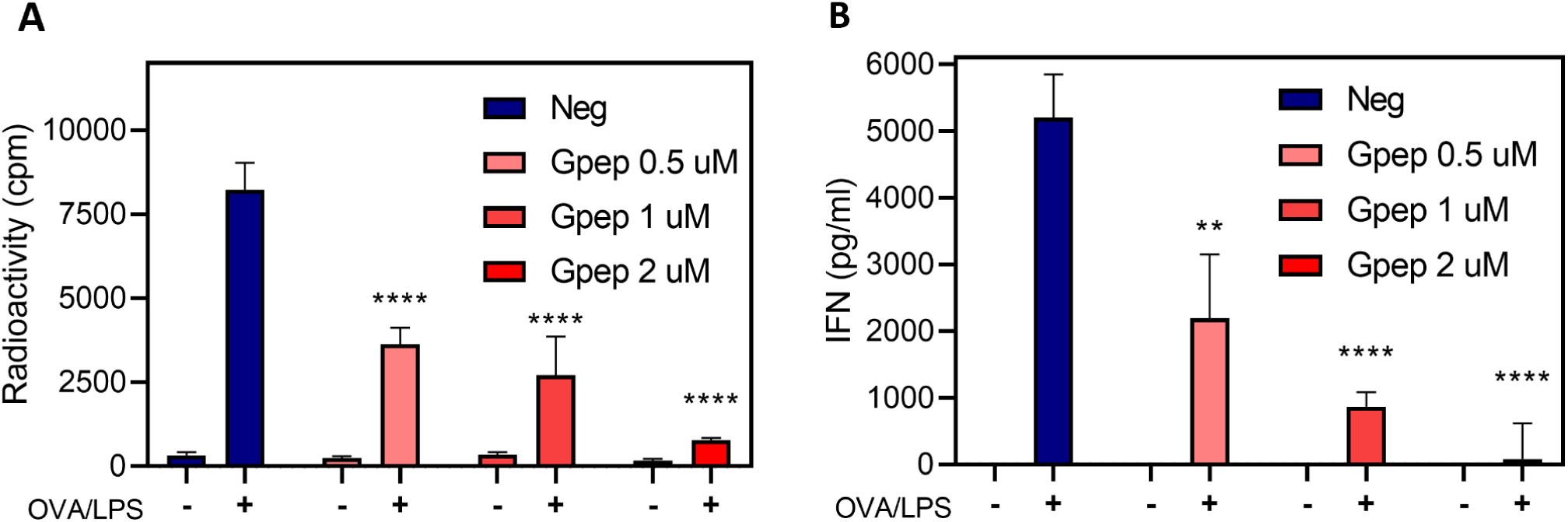
Gpep inhibits splenocyte activation in a dose dependent manner. (A) Splenocytes were recovered from the spleens of OT-II mice, red blood cells were lysed, and cells were incubated for 18 hours at 37°C in the absence (-) or presence (+) of activation stimuli (500 ng/mL LPS and 0.1 μg/mL OVA peptide), with Gpep at 0, 0.5, 1, or 2 μM concentrations. At this time point, IFN-γ levels in the supernatants were quantified by indirect ELISA. (B) Cells were incubated for 48 hours in the presence of the stimuli and [³H]-methylthymidine. [³H]-methylthymidine incorporation was measured to assess cell proliferation. Bars represent the geometric mean ± SD of radioactivity (CPM) or IFN-γ concentration (pg/mL). *p<0.05, **p<0.01, ***p<0.001, ****p<0.0001 (ANOVA, all comparisons to Neg). Depicted assays are representative of at least two independent experiments.

The immunosuppressive activity of Gpep extends to human neutrophils, a cell type implicated in both protective and pathological responses during RSV infection. Functional assays revealed that Gpep impaired neutrophil chemotaxis toward fMLP, suppressed LPS-induced CD11b upregulation, and reduced fMLP-induced reactive oxygen species (ROS) production (**Fig. 7A-C** and **Fig. S3A-B**). Additionally, Gpep inhibited PMA-induced myeloperoxidase (MPO) release and neutrophil extracellular trap (NET) formation, as measured by extracellular dsDNA quantification (**Fig. 7D**-**6E**). Importantly, these inhibitory effects occurred without compromising neutrophil viability (**Fig. Sup 3C**), confirming that Gpep specifically attenuates neutrophil activation rather than inducing cytotoxicity.

**Figure 7.**
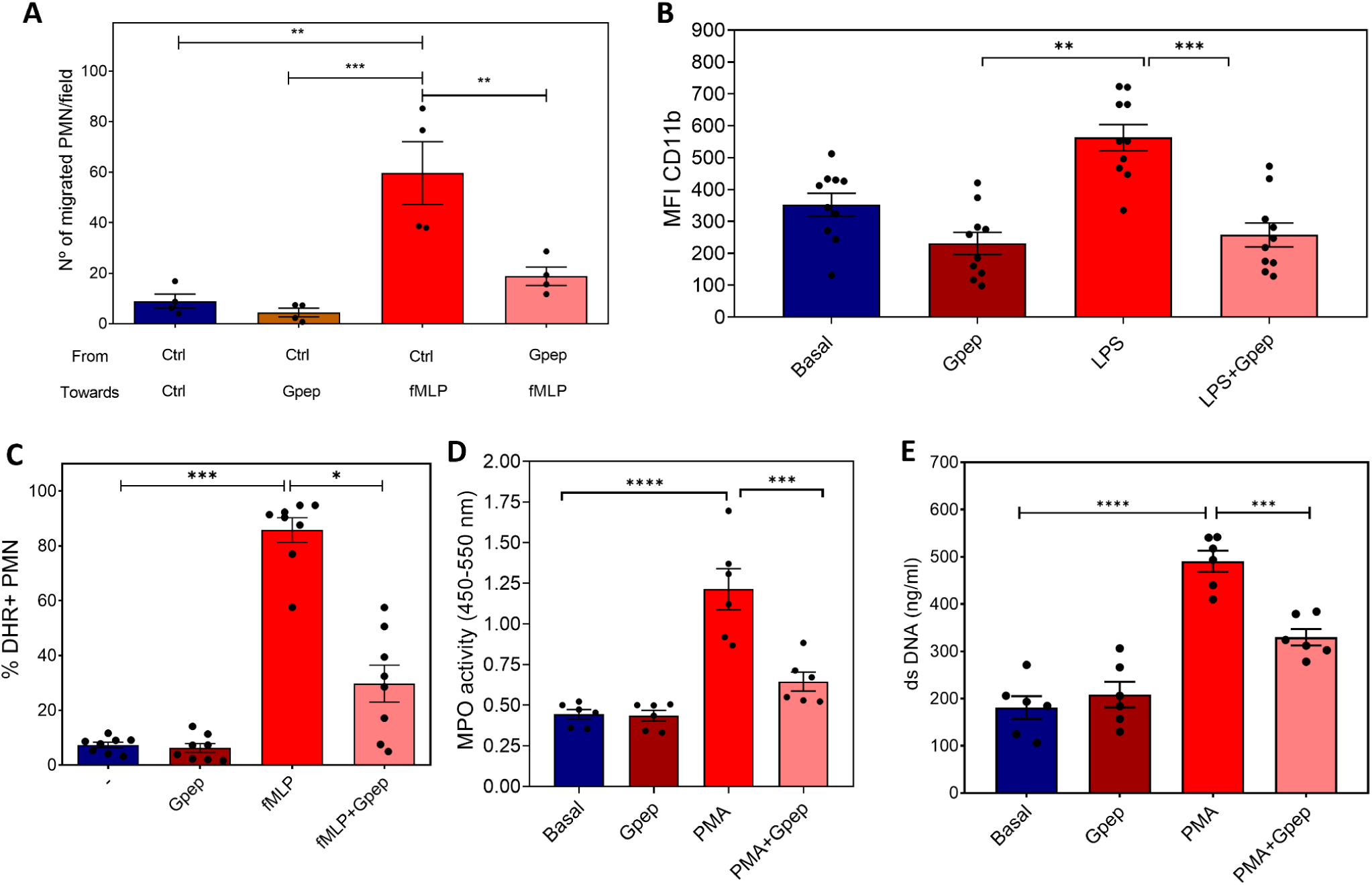
Gpep inhibits human neutrophil activation in vitro. (A) Neutrophil migration was evaluated using a modified Boyden chamber assay in the presence or absence of Gpep 1 uM. Migrated cells were quantified after 30 min and results expressed as the percentage of input cells. (B) Surface CD11b expression. Neutrophils were incubated for 30 min with Gpep 1 uM or vehicle and stimulated with fMLP (10^-7^ M). The mean fluorescence intensity (MFI) of CD11b was determined by flow cytometry. (C) Reactive oxygen species (ROS) production. Neutrophils were incubated for 30 min with Gpep or vehicle and stimulated with fMLP (10^-7^ M). ROS generation was quantified by flow cytometry using the DHR probe, and results are expressed as the percentage of DHR⁺ cells. (D) Myeloperoxidase (MPO) release was quantified in cell-free supernatants collected after 2 h of stimulation, measured spectrophotometrically, indicating primary granule mobilization. (E) Neutrophil extracellular trap (NET) formation. Extracellular DNA was quantified after 3 h stimulation, as an indicator of NET release. In all cases, results are expressed as mean ± SEM from at least four donors. *p<0.05, **p<0.01, ***p<0.001, ****p<0.0001 by ANOVA, all comparisons to Control + (LPS, fMLP, PMA)

Notably, the immunomodulatory activity of Gpep was dependent on its oligomerization state. Concentrated Gpep (200 μM vs 20 μM, **Fig. 2b**) that results in an oligomerized form of Gpep, displayed a marked loss of immunomodulatory function within physiologically relevant concentrations (**Fig. Sup 4**), paralleling the lack of immunosuppressive activity observed for membrane-bound G (mG) in inactivated RSV. These findings suggest that the oligomeric form of Gpep does not engage in immunomodulation, likely due to conformational constraints that prevent interaction with immune receptors and/or disruption of the CX3C motif. Thus, only the monomeric, native-like form of Gpep is capable of exerting immunosuppressive effects, highlighting a potential functional divergence between soluble and membrane-bound states of the RSV G protein.

### 3. Divergent Kinetics of Anti-G and Anti-F IgG Antibodies During Early Pediatric RSV Exposure

To elucidate the humoral immune response dynamics following primary and early exposures to RSV, and to specifically assess the role of the immunomodulatory CX3C motif within the G protein, we quantified IgG antibody titers against the two principal viral antigens, F and G, in sera from a cohort of healthy, ambulatory pediatric patients aged 0 to 72 months (see **Fig. 8**). Stratification by age allowed us to distinguish between groups predominantly protected by maternally derived antibodies (0–6 months) and those where endogenous, adaptive immunity increasingly contributes (6-12, 12–24 and 24–72 months).

**Figure 8.**
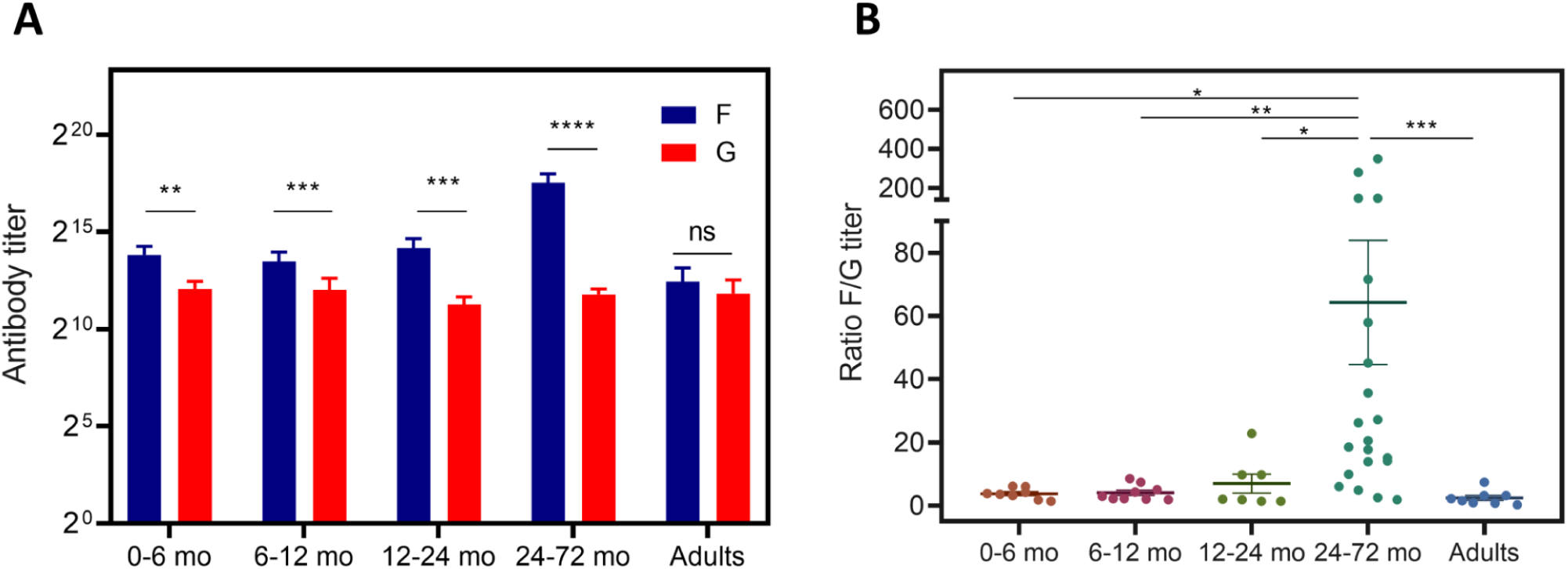
Uncoupling of anti-G and anti-F responses during early exposures to human RSV. (A) Anti-F and anti-G antibody titers across different age groups are shown as bar graphs representing mean ± SEM. Each age group includes paired measurements of anti-F and anti-G titers from individual donors. Paired data were compared using the Wilcoxon signed-rank test. *p < 0.05, **p < 0.01, ***p < 0.001, ****p < 0.0001, ns = not significant. (B) Ratio of anti-F to anti-G titers (F/G ratio) represented as a dot plot, where each dot corresponds to an individual donor. Data were analyzed by Kruskal–Wallis test followed by Dunn’s post-hoc test. The horizontal lines indicate the mean value and SEM. *p < 0.05, **p < 0.01, ***p < 0.001, ****p < 0.0001, ns = not significant.

Analysis of antigen-specific antibody titers revealed a clear divergence in the kinetics of anti-F and anti-G antibody responses across pediatric and adult age groups. In all pediatric age groups anti-F IgG were more abundant than anti-G, contrasting with adults where no significant differences were found between antigens (**Fig. 8A**). In the youngest infants this difference was slighter, reflecting a greater contribution of maternal transfer. As patient age increased, the disparity between antigen titers became more pronounced, indicating a predominance of anti-F IgG as active immunity begins to emerge. Notably, in the 24-72 months old cohort, a marked divergence was observed, with anti-F titer greatly exceeding those against G. The F/G IgG titer ratio in this group was distinct from all other age groups, underscoring a delayed or less robust anti-G antibody response relative to anti-F during the period of active immune maturation (**Fig. 8B**). These findings collectively suggest that while anti-G antibodies are efficiently transferred, the development of anti-G responses, potentially influenced by the structural and immunomodulatory properties of the G protein’s CCD, lags behind, becoming quantitatively similar to anti-F only after repeated antigenic encounters.

## Discussion

This study provides a comprehensive characterization of the RSV G protein central conserved domain (CCD), elucidating its biochemical properties, immunomodulatory capacity, and displaying its potential impact on humoral immunity in pediatric populations, with direct implications for vaccine design. The principal findings demonstrate that the CCD’s four cysteine residues form a putatively native CX3C chemokine-like motif at low concentrations, which is structurally stabilized by disulfide bonds and capable of binding the CX3CR1 receptor expressed on human airway epithelial cells (hEAC) and diverse immune cell subsets^57^. Functional assays revealed that the CCD, when expressed as a recombinant peptide (Gpep), selectively suppresses activation of dendritic cells (DCs), monocytes, T cells, and neutrophils in-vitro, without inducing cytotoxicity. To the best of our knowledge, this is the first report showing the effect of RSV G on neutrophils. Neutrophil recruitment is an early event following RSV infection acting as a first line of defense in viral clearance and also promoting adaptive immunity^58^. Treatment with the Gpep significantly attenuated neutrophil inflammatory responses in-vitro, as evidenced by impaired chemotaxis, reduced generation of reactive oxygen species (ROS), diminished release of myeloperoxidase (MPO), and suppression of neutrophil extracellular trap (NET) formation. Thus, collectively our results show that the G peptide is able to modulate both the innate and adaptive immune responses. Notably, this immunosuppressive activity is contingent upon the peptide’s oligomerization state and concentration, with monomeric Gpep displaying robust function, while oligomeric forms and/or membrane-bound G (mG) lack immunomodulatory effects.

The versatility of CCD disulfide bonds emerged as a key determinant of structural and functional outcomes. Oxidative folding studies under varied redox conditions demonstrated that Gpep rapidly attained its native state, with minimal accumulation of folding intermediates sensitive to disulfide reshuffling. At higher concentrations, Gpep undergoes intermolecular disulfide bonding, suggesting that oligomerization of membrane-bound G may be similarly mediated, although the precise mechanism remains unresolved^23,59,60^. Molecular dynamics simulations, size-exclusion chromatography, and fluorescence spectroscopy collectively indicate that reduced Gpep adopts a compact, metastable fold, while the native, disulfide-bonded peptide is more extended in solution, likely due to unstructured N-terminal regions. These findings highlight the concentration-dependent conformational plasticity of the CCD, which may underlie distinct functional profiles in the soluble and membrane-bound forms of the G protein.

Structurally, the distinction between sG and mG forms is consequential for immunomodulation. The data show that only the monomeric, soluble Gpep retains the ability to attenuate innate immune activation, suppressing key responses in dendritic cells and neutrophils, while oligomeric CCD and inactivated RSV, where mG predominates, fail to elicit these effects. This implicates sG as the principal mediator of RSV-driven immunosuppression via the CX3C motif, with oligomerization and membrane association possibly imposing conformational constraints that may preclude receptor engagement and functional modulation. Reversible formation and reduction of disulfide bonds in redox-sensitive motifs within viral ectodomains can induce structural rearrangements, resulting in functional changes. For instance, redox regulated mechanisms for viral entry and membrane fusion have been described for HIV Gp120/Gp41 protein^61,62^. Allosteric disulphide bridges have also been described in multiple extracellular molecules such as integrins, where the cysteine residues act as redox switches modulating structure-function relationships^63^. In the case of RSV CCD, the differential activities observed underscore the importance of CCD structure and presentation in shaping the host immune response.

Observations in pediatric cohorts further delineate the functional consequences of CCD immunomodulation on humoral immunity. Analysis of anti-F and anti-G IgG antibody titers in children aged 0–72 months revealed a pronounced bias toward anti-F responses during early exposures, with anti-G titers lagging, particularly in the 24–72 month group. This temporal dissociation was not observed in adults, where antibody levels against both antigens equilibrate, nor in infants within the window of maternal antibody persistence, who display ratios similar to adults. The delayed rise of anti-G antibodies in children may reflect the immunosuppressive activity of the CCD, differences in antigen processing, or selective modulation of adaptive responses during initial RSV encounters. Notably, these findings align with previous reports that associate lower anti-G titers with increased disease severity^64^ and highlight the potential for sG-mediated immunosuppression to influence the trajectory of immune maturation and clinical outcomes^21,24^. Whether the secreted sG protein shapes the humoral immune response during early RSV infections remains an open question that requires further research.

Collectively, these results have significant implications for RSV vaccine design, particularly in pediatric populations at heightened risk for severe disease. The capacity of the G protein CCD to suppress PAMP-mediated activation of innate immunity during the early stages of infection, when sG concentrations are highest, may dampen the activation of antigen-presenting cells and skew the development of protective immune memory. Coupled with additional viral mechanisms of immune evasion, such as IFN suppression by NS1 and NS2 proteins^9,10^, RSV employs a multifaceted strategy to silence host defenses. Therefore, effective vaccine strategies must not only target key antigenic determinants but also counteract CCD-mediated immunosuppression, ensuring robust activation of both innate and adaptive immunity in young children. Elucidating the mechanisms by which the CX3C motif modulates immune responses will be critical for preventing severe RSV disease and establishing durable protective immunity.

## Associated Content

Supporting Information available:

Includes:

**Figure S1.** Purification workflow of RSV G peptide (Gpep), including affinity, ion-exchange, and size-exclusion chromatography, and final purity assessment.

**Figure S2.** Immunogenicity and immunoreactivity of Gpep in human sera and in mice, including ELISA and Western blot analyses.

**Figure S3.** Cytometry data of PMN activation, including ROS generation, CD11b expression, and cell viability.

**Figure S4.** Loss of Gpep inhibitory activity upon oligomerization, evaluated by IFN-γ production and neutrophil ROS assays.

**Figure S5.** Representative ELISA titration curves stratified by age group.

## Acknowledgments

This work was funded by the National Agency for the Promotion of Research,Technological Development and Innovation (ANPCyT) ( PICT 2019 −03029, PICT 2020-00067.) and by the national scientific and technical research council (CONICET), (PIP 2021-2023) SAE, DAP MVT, MTC and GCF are members of the scientific and technological research career of the national scientific and technical research council (CONICET), Argentina.

