## supplementary material for "Unraveling the Dual Immunomodulatory and Immunogenic Roles of the Central Conserved Cysteine-Rich Region in Respiratory Syncytial Virus G Protein"

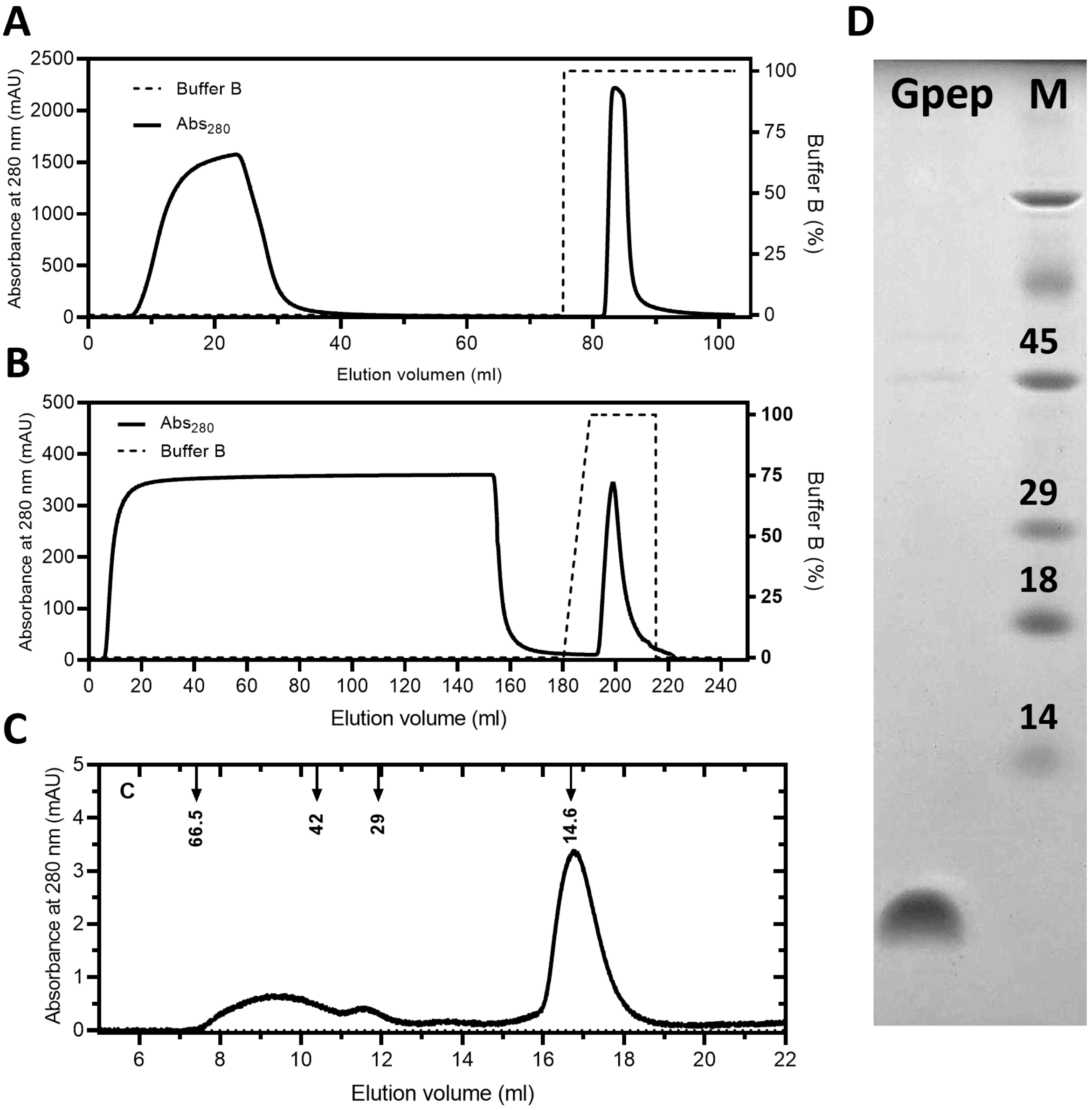


**Figure Sup 1. Purification of RSV G peptide**. (A) Amylose affinity chromatography of the MBP-Gpep fusion protein. The fusion protein was bound to the amylose resin and eluted with 20 mM maltose in buffer A (20 mM Tris-HCl, 200 mM NaCl, pH 7.6), producing a distinct elution peak corresponding to MBP-Gpep. (B) Cation exchange chromatography (Capto S) of TEV cleaved Gpep, exploiting its high isoelectric point (pI = 9.5) relative to MBP (pI = 5.1), eluted with 500 mM NaCl in buffer A (phosphate 50 mM pH 7.6). (C) Size exclusion chromatography (Superdex 75 10/300) in PBS of Gpep. (D) Tris-Tricine SDS-PAGE analysis of the final preparation, confirming ~95% purity of Gpep.Lane 1: Purified Gpep; Lane 2: molecular weight marker (kDa)

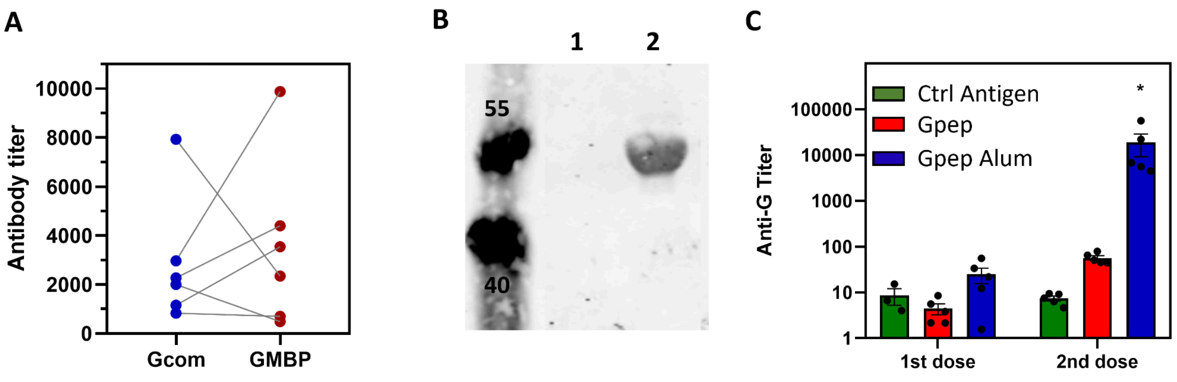


**Figure Sup 2. Gpep immunogenicity and immunoreactivity with RSV-specific sera and Anti-G protein antibodies.** (A) Adult sera recognize the Gpep-MBP fusion protein with titers comparable to those against the full-length G protein in an ELISA assay. Paired serum samples were analyzed, and antibody titers were compared using a Wilcoxon matched-pairs signed rank test. No statistically significant difference was observed between anti-G and anti-Gpep-MBP responses ($p > 0.5$). (B) A commercial anti-G antibody (Catalog 40626-T62, SinoBiological) recognizes the G-MBP fusion protein (lane 2) but not MBP alone (lane 1) in Western-blot. (C) Sera from BALB/C mice (n=5) immunized with Gpep adjuvanted with alum recognize the full-length G protein in an ELISA assay. Bars represent the geometric mean ± SEM of anti-G titer. *p<0.05, **p<0.01, ***p<0.001, ****p<0.0001 (ANOVA, all compared to Control -).

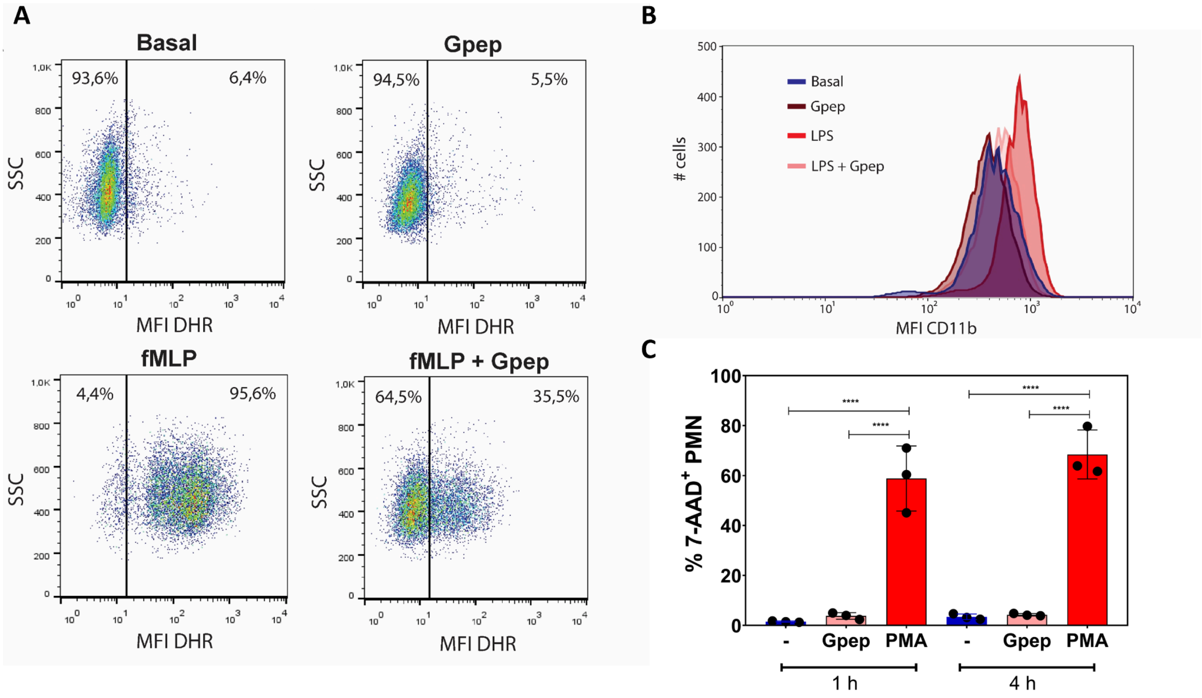


**Figure Sup 3: PMN additional plots.** (A) Dot-plots of DHR fluorescence versus side scatter (SSC) illustrate reactive oxygen species (ROS) generation in neutrophils under different conditions, corresponding to the data shown in Figure 7C. (B) Representative histograms of CD11b surface expression on neutrophils after stimulation, corresponding to Figure 7B. Fluorescence intensity reflects the level of activation marker upregulation. (C) Neutrophils ($5x10^5$) were incubated with Gpep or a positive control of death (PMA, 100 nM) for 1 or 4 h at 37 °C in 5 % $CO_2$. After the incubation period, cells were washed and incubated with 7-Amino-Actimycin D (7-AAD; BD Biosciences, 10 µg/ml) for 10 min on ice in the darkness. Immediately after, cell viability was determined by flow cytometry, since 7-AAD is excluded by viable cells but can penetrate cell membranes of dying or dead cells. Only PMA stimulation results in a detectable increase in $7\text{-}AAD^+$ cells (%$7\text{-}AAD^+$ PMN), indicating minimal apoptosis under control and Gpep conditions.

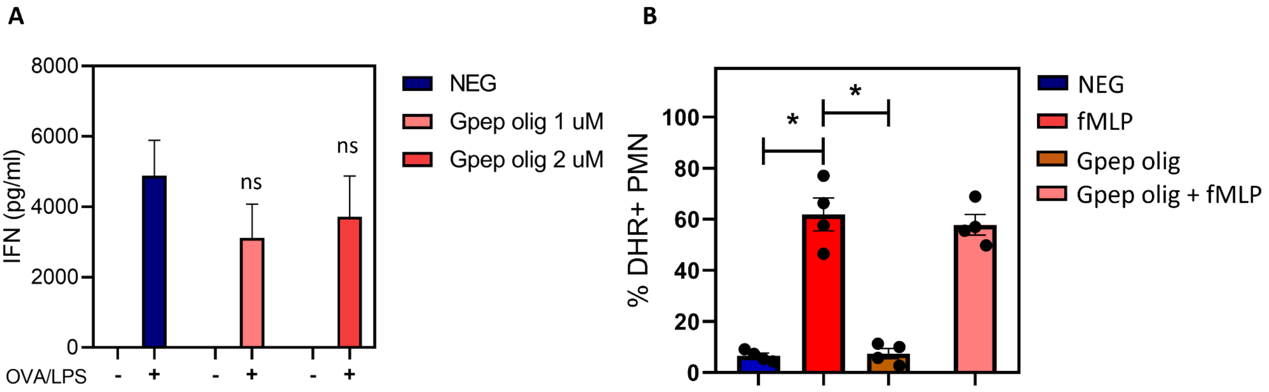


**Figure Sup 4: Gpep's inhibitory activity is lost upon oligomerization.** (A) Splenocytes were recovered from the spleens of OT-II mice, red blood cells were lysed, and cells were incubated for 18 hours at 37°C in the presence or absence of activation stimuli (500 ng/mL LPS and 0.1 µg/mL OVA peptide), with Gpep at 1 or 2 µM. Control − refers to cells without any stimulus. At this time, IFN-γ levels in the supernatants were quantified by indirect ELISA. Bars represent the geometric mean ± SD of IFN-γ concentration (pg/mL). No significant differences were observed in any comparison with the negative control. (B) Reactive oxygen species (ROS) production. Neutrophils were incubated for 30 min with Gpep 1 uM or vehicle and stimulated with fMLP ($10^{-7}$ M). ROS generation was quantified by flow cytometry using the DHR probe, and results are expressed as the percentage of $DHR^{+}$ cells. Statistical analysis was performed using one-way ANOVA.

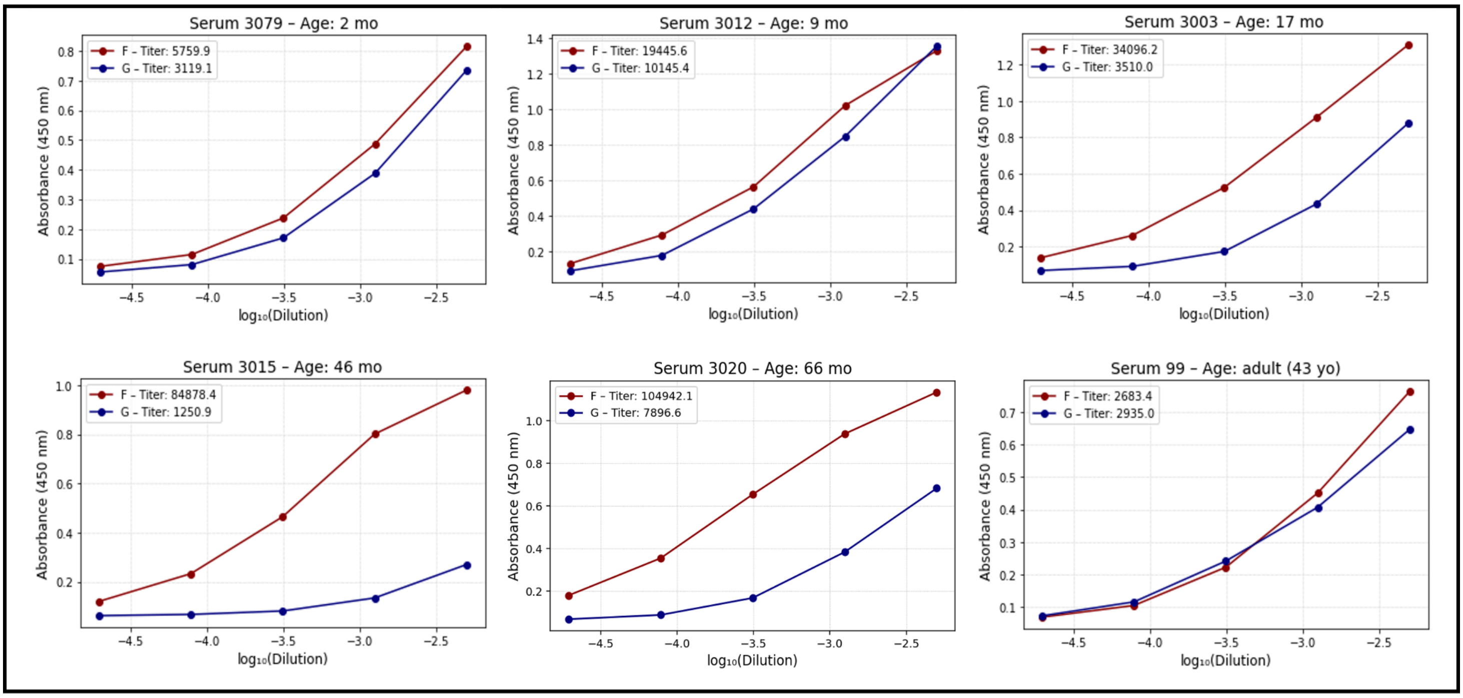


**Figure Sup 5: Representative ELISA titration curves of sera from different age groups.** Sera were serially diluted and tested for reactivity against RSV F and G antigens in the same plate. The curves illustrate the dilution-dependent signal decay and the relative differences in antibody titers among age groups.
